# Contractility, Differential Tension and Membrane Removal direct zebrafish Epiboly Biomechanics

**DOI:** 10.1101/113282

**Authors:** Maria Marsal, Enrique Martin-Blanco

## Abstract

Precise tissue remodeling during development is essential for shaping embryos and for optimal organ function. Epiboly is an early gastrulation event by which the blastoderm expands around the yolk to engulf it. Three different layers are involved, an epithelial layer (the enveloping layer, EVL), the embryo proper, constituted by the deep cells (DCs), and the yolk cell. Although teleost epiboly has been studied for many years, a clear understanding of its mechanics was still missing. Here we present new information on the cellular, molecular and mechanical elements involved in epiboly that, together with some other recent data and upon comparison with previous biomechanical models, lets conclude that the expansion of the epithelia is passive and driven by cortical contraction and membrane removal in the adjacent layer, the External Yolk Syncytial Layer (E-YSL). The isotropic actomyosin contraction of the E-YSL generates an anisotropic stress pattern and a directional net movement as a result of the differences in the deformation response of two opposites adjacent domains (the EVL and the Yolk Cytoplasmic Layer - YCL). Contractility is accompanied by the local formation of membrane folds and the membrane removal by Rab5ab dependent macropinocytosis. The increase in area of the epithelia during the expansion is achieved by cell-shape changes (flattening) responding to spherical geometrical cues. The counterbalance between the geometry of the embryo and forces dissipation is therefore essential for epiboly global coordination.

## MAIN TEXT

To fulfill organ functional optimization, during development cells within tissues change positions and shapes coordinating with each other through time. Morphogenetic processes demand cells motion and motility depend on forces. Forces act by directing mechanical action and reaction processes within tissues and amongst cells.

While our knowledge of the signaling networks and structural elements involved in morphogenetic processes is substantial, very little is known on the mechanical control of morphogenesis. Both, mechanosensation and mechanotransduction are largely unexplored throughout development. Further, the influence that tissues mechanical properties may have modulating cellular responses remains undefined.

Epiboly in Teleosts constitutes one of the best models to study the mechanical control of morphogenesis. The first detailed descriptions of epiboly come from *Fundulus* ^1^. Epiboly consist on the coordinated expansion of three adjacent tissues, the external Enveloping Layer (EVL), a cohesive epithelium, the internal Yolk Syncytial Layer (YSL), a syncytial layer beneath, and the Deep Cells (DCs), the embryo proper cells, sandwiched in between. These tissues expansion proceeds from one of the poles of the embryo to the opposite (in an animal (A) to vegetal (V) direction). Expansion occurs at the expense of the Yolk Cytoplasmic Layer (YCL) at the surface of the yolk cell. This description of epiboly is also valid for the zebrafish. Importantly, all the events involved in epiboly take place simultaneously and must be interpreted in the context of the whole embryo dynamical equilibrium (volume conservation and invariable spherical shape).

### Geometrical and non-geometrical constrains during epiboly progression

The spherical geometry of teleosts embryos imposes two different scenarios during epiboly progression. As the EVL moves forward, the perimeter of its margin increases until it reaches the equator, while the yolk domes, yolk granules initiate toroidal movements and an actin-rich domain at the yolk surface adjacent to the EVL, the External-YSL (E-YSL) is singled out. In a second stage, the perimeter of the advancing front progressively decreases up to closing at the V pole and yolk granule movements, which halt as the EVL surpass the equator, reinitiate ^2^ Remarkably, epiboly does not seem to rely in cell proliferation ^3–9^; and the inhibition of cell divisions just shows very minor epiboly phenotypes ^10^. Cell divisions orientation does not seem to affect directional tissue growth either and certainly does not influence the anisotropic tissue tension of the EVL or promotes tissue spreading ^10^. Overall, the surface area of the whole EVL increases around 3,3 times (Figure 1A) and this increase directly correlates with a largely uniform reduction (flattening) of EVL cells height (Figure 1B).

**Figure 1.**
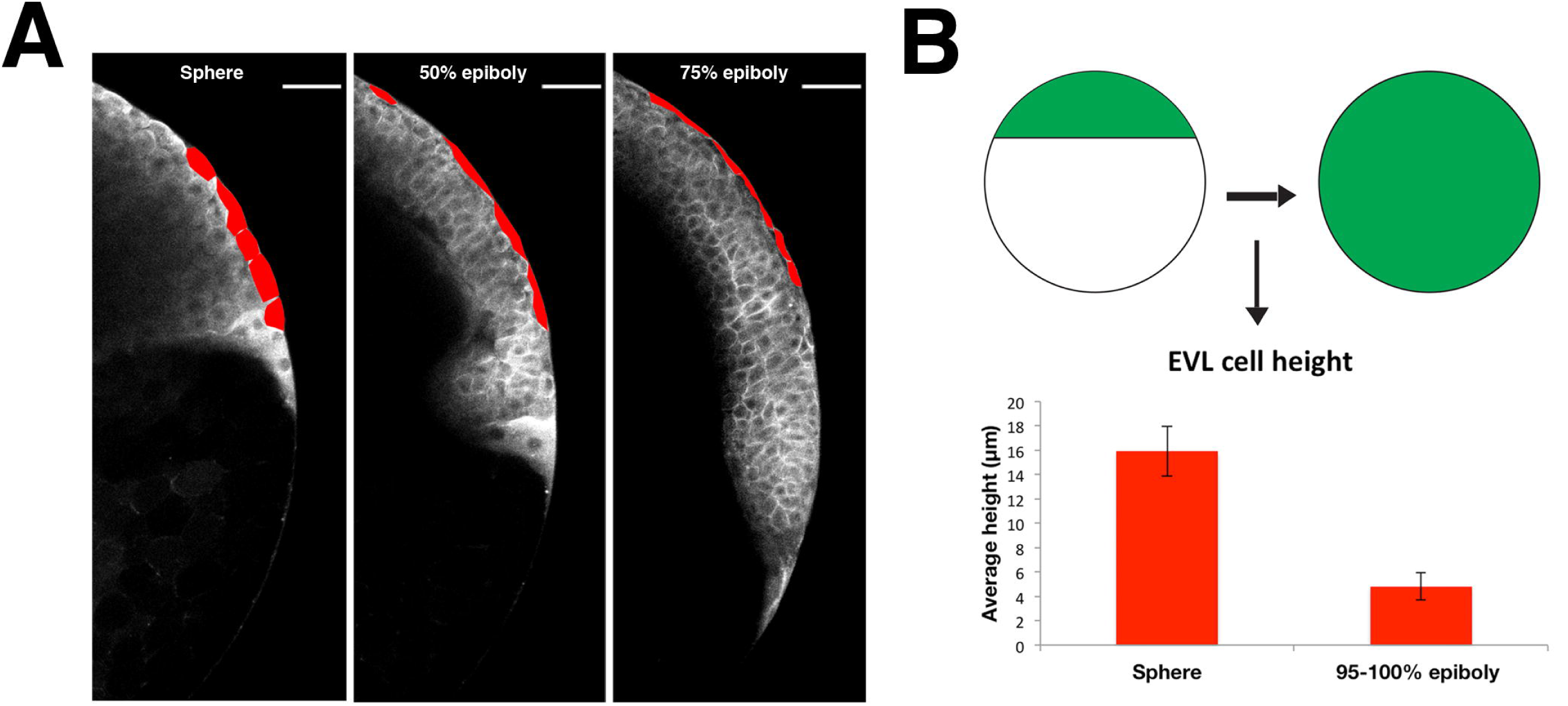
EVL cells flattening accounts for most of its expansion throughout epiboly. **A)** Snapshots of a two-photon microscopy time-lapse movie at the medial plane of a membrane labeled transgenic embryo [Tg (β-actin:mGFP)]. The lateral outlines of distinct EVL cells are marked in red. Scale bar, 50 μm. **B)** Geometrical estimation of the increment in surface area of the EVL during epiboly. The area of the surface of the sector covered by the EVL on an imaginary sphere increases 3.3 times from epiboly onset to closure. On the other hand, average EVL cells height at sphere (15,9 ± 2,1 μm) and epiboly closure (4,8 ± 1,1 μm) show also a reduction of 3,3 times. Error bars - Standard Deviation, n=16 for each condition, data from 2 embryos.

An obvious difference between the early and late stages of epiboly progression is the morphology of the EVL cells at the margin. Initially, these cells are misaligned and display an extensive set of apical filopodia; after surpassing the equator, however, they stop emitting protrusions and altogether are brought into line ^11–13^. Cell morphologies are thus reflecting geometrically imposed restrictions (the shift from an expanding to a shrinking front).

Conversely, geometry does not influence the dynamics of the actin-rich E-YSL or the yolk doming. The E-YSL, which initially covers a wide domain adjacent to the EVL, narrows down up to a time when its width gets stabilized. Remarkably, neither in *Fundulus* ^1, 5^ nor in zebrafish E-YSL shrinking correlates with geometrical restrictions. The narrowing of the E-YSL in *Fundulus* comes to an end before the EVL reaches the equator, at around 42% epiboly ^1, 5^, while, in zebrafish, it ceases after the EVL overpasses the equator, at around 60% epiboly. The actual yolk doming does not follow the expansion of the EVL either. Doming depth is proportional to the area occupied by the blastoderm at the onset of epiboly; 18-19% in *Fundulus* (Stage 13) and 30% in zebrafish.

Some evidences from *Fundulus* suggest that the narrowing of the E-YSL and the doming of the yolk could be mechanically linked ^1, 5^. In response to experimentally detaching the EVL-E-YSL junction before the onset of epiboly (Stage 11), the free denuded yolk immediately domes, while the narrowing of the E-YSL proceeds ahead of time ^14^ Both doming and E-YSL narrowing in these denuded eggs are faster than in intact embryos ^1, 5^, suggesting that the blastoderm cap conveys an intrinsic resistance to epiboly progression. Moreover, in the zebrafish, the narrowing of the E-YSL temporally associates with the ingression of internal yolk syncytial nuclei ^15^ to constitute the Internal YSL (I-YSL) ^16, 17^, which also strictly correlates with the doming of the yolk. This, itself, could be modulating the extent and timing of E-YSL condensation (Figure 2A).

**Figure 2.**
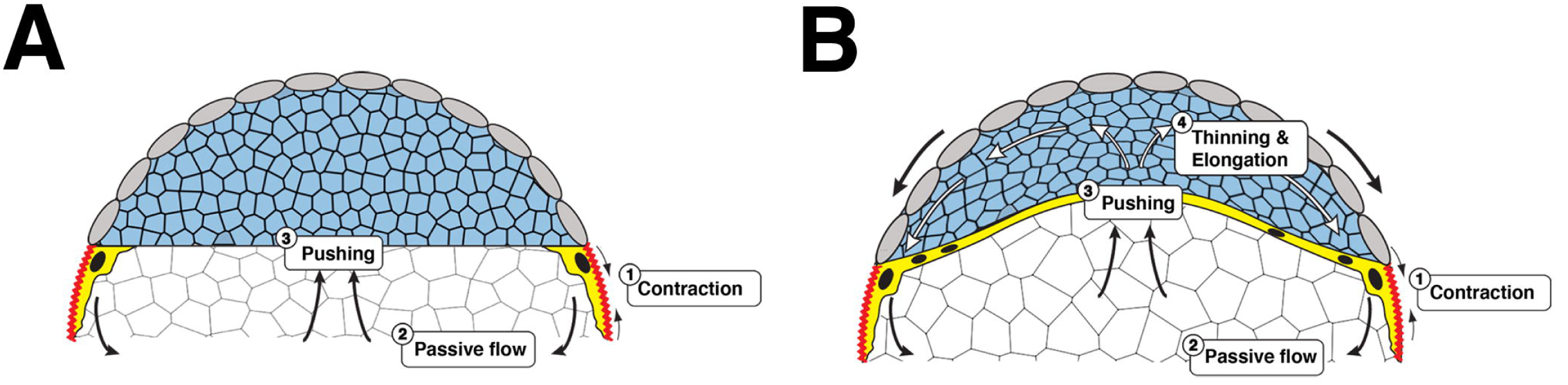
An early mechanical feedback loop for doming and EVL spreading. A and B) Schematic representation of the cross-section of an embryo just after sphere stage (A) and during doming (B). The initial contraction of the E-YSL would induce the passive flow of yolk granules, which, in sequence, will push the DC basal margin. When the forces generated by the passive flows will overcome the resistance of the blastoderm, the DCs will get displaced upwards. As the DCs are spatially constrained by the EVL and this is tightly attached to the yolk at the surface (EVL-E-YSL attachment), the DCs’ layer will radially intercalate and elongate in the A-V direction. In turn, this would generate a frictional force in the EVL basal side and an extra impulse into the EVL-E-YSL attachment, favoring E-YSL narrowing and EVL epiboly through a mechanical positive feedback loop.

### An early mechanical feed back loop for yolk doming

The contractile capability of the E-YSL appears to be instrumental for epiboly initiation. Considering that both, embryo volume and shape are constant, its active shrinkage will pull from the adjacent EVL and induce an overall redistribution of the yolk incompressible internal elements leading to the passive flows of yolk granules. At the geometrical centre of the embryo the yolk granules would flow animalward pushing the basal side of DCs layer.

We hypothesized that if the stress generated by the yolk granules in the DC’s basal side overcomes these cells resistance, doming would take place. This movement, as the DCs are spatially constrained by the EVL, would result into an additional mechanical input accelerating the vegetalward progression of the EVL/E-YSL border. The radial intercalation (lengthening) of DCs during doming ^18, 19^ may also reinforce this positive feedback, pushing the E-YSL and strengthening its contraction. The narrowing of the surface area occupied by the yolk nuclei (E-YSL) and the formation of the I-YSL ^15^ would simply be indirect consequences of the initial E-YSL contraction passively transmitted via yolk granules flows and yolk doming (Figure 2B).

### Differential tension model of epiboly progression: Stress and Power

The contractile activity of the E-YSL during epiboly generates stress at its opposite adjacent structures, the EVL and the YCL. By experimentally increasing cortex contraction at the E-YSL (wound healing response), the EVL cells respond by passively deforming and expanding by cell flattening. On the opposite side, the thin YCL cannot stretch in response to contraction.

As epiboly progress, the YCL gets incorporated into the E-YSL, where the yolk membrane is locally removed. We propose that epiboly is just the consequence of the E-YSL active contractility transmitted to the YCL and of yolk cell membrane turnover (Figure 3A and 3B) ^2^

**Figure 3.**
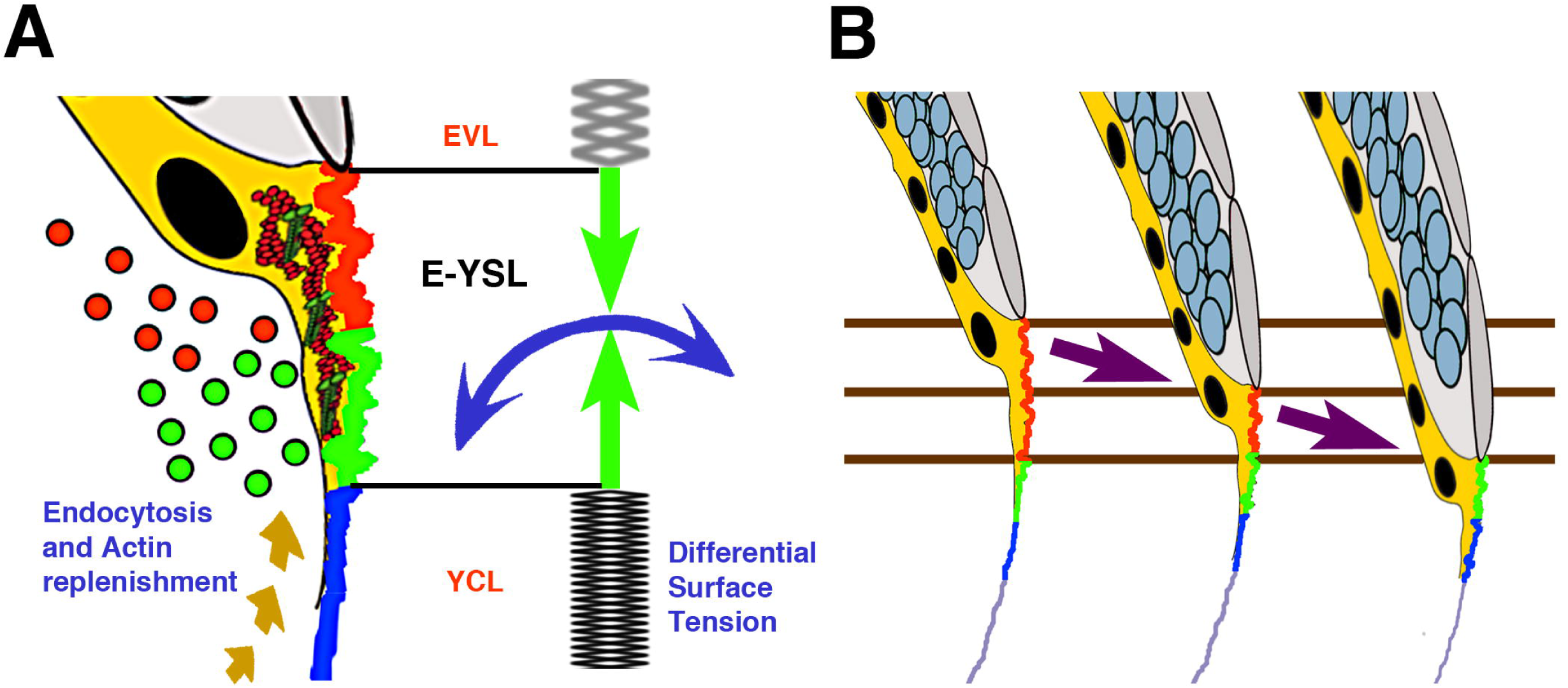
Differential tension model of epiboly progression. **A)** The movement of the EVL and DCs towards the vegetal pole demands the removal of the yolk cell surface at the E-YSL. Different zones on the surface of the E-YSL and endocytosed vesicles (dots) are color coded in red and green. Membrane removal associates to contraction of the E-YSL surface and the recruitment of actin and myosin (beige dashed arrow) from vegetally located pools. Actin and myosin are diagrammatically illustrated in red and green within the YSL (yellow). The E-YSL contracts (CC - blue and AV - green) and pulls both adjacent domains, the EVL and the YCL. The imbalance of stiffness between the EVL (elastic) and the yolk cell surface (YCL - rigid) account for epiboly progression. The EVL expands (passive cell flattening) while the YCL cannot deform. **B)** Three chronological time points of epiboly progression are shown. The EVL (grey) and the DCs (pale blue) move towards the V pole while the E-YSL (red and green) undergoes a rapid turnover progressively incorporating the smooth YCL (blue).

This differential tension model is supported by multiple observations including: the dynamic pattern of stresses at the yolk cell surface showing a graded animal-vegetal (AV) distribution; the yolk membrane kinematics pointing to a static YCL; and the response of the EVL to the contraction at the adjacent E-YSL, deforming more than the yolk cell surface. The different responses of the EVL and the YCL (with different mechanical properties, elastic and rigid) to the isotropic cortex contraction of the E-YSL will result into the movement of the EVL in the V direction. E-YSL endocytosis and the progressive conversion of the adjacent YCL into E-YSL would constitute necessary and essential factors.

Power density dynamics inferred by Hydrodynamic Regression (HR) clearly point to an unmistakable source of active forces at the E-YSL from 50 % epiboly onwards ^2^ However, no positive mechanical activity at this region was detected before the EVL reached the equator. If, indeed, the positive feedback loop suggested above contributes to the first phase of epiboly, just small increments in the contraction of the E-YSL could be sufficient to trigger the expansion of the EVL and the mesoscopical HR analyses may not be sensitive enough to detect it. During the second phase of epiboly (after 50 %), the positive feedback loop between the three layers would become negligible (no significant doming is observed in this period) and HR, which is based on yolk granules flow patterns, detects with high confidence the increased positive power of the E-YSL (contractility) and the resistance to stress of the EVL.

### Local sources of power: membrane removal and cortex contractility

EVL cells do not slide over the yolk cell and, therefore, to allow for epiboly progression, the yolk cell membrane should withdraw. Membrane removal during epiboly occurs within the E-YSL and is dependent on the activity of the small GTPase Rab5ab ^20^. Remarkably, the knockdown of Rab5ab function results in a delay in doming and in the formation of a free space between the DCs and the yolk (compare Figure 4A and 4B). Thus, the endocytosis of the yolk cell membrane is an essential element for the doming of the blastoderm. Reducing Rab5ab activity and hence, E-YSL endocytosis would result in the mechanical impairment of the dissipation of stress into the yolk and, as a consequence, in doming failure.

**Figure 4.**
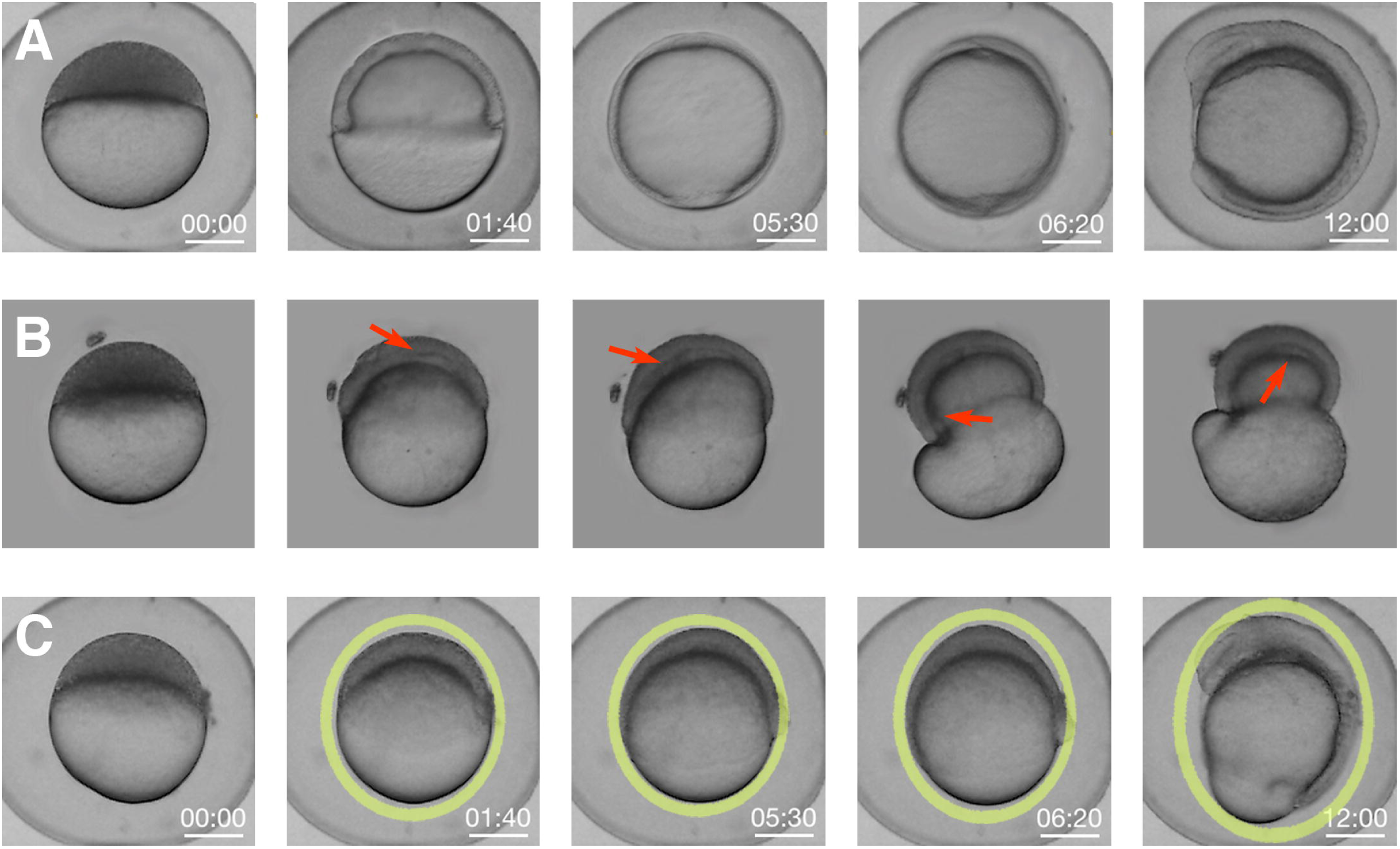
Knockdown of Rab5ab function results in tensional unbalance. **A)** Macroscopic bright field images of sibling controls embryos imaged within their chorion at different stages. Scale bar 250 μm. **B**) High (8 ng) dose Rab5ab YMO embedded in agarose under space constrain showing epiboly delay (remaining at 40 % epiboly when control siblings have already closed) and a gaps/detachments between the DCs and the yolk cell (red arrows). **C)** Rab5ab YMOs not subjected to mechanical constrains imaged within their chorion lose their spherical shape (oval yellow rings).

In wild type conditions the embryo sustains a spherical shape during the whole expansion. However, in Rab5ab yolk-depleted embryos the spherical shape is lost (compare Figure 4A and 4C). This shape alteration has also been observed upon impairing EVL progression after interference in other genes’ functions ^21, 22^ Mildly compromised embryos (Rab5ab medium dose MOs) will elongate in the AV direction compensating stresses and balancing disturbed pushing forces. For high dose MOs with extremely impaired endocytosis and nearly no V EVL displacement, no force compensation will take place, the embryos will not change shape and the yolk will burst.

The overall shape distortion of Rab5ab embryos is consequence of the uncoupling between EVL expansion and DCs progression. DC’s are unable to progress in the V direction as their movements are impeded by the slow moving EVL-E-YSL attachment. In these embryos, the cortical power density is strongly diminished ^20^ while the pushing forces for DCs ingression appear to be sustained. The equilibrium of forces between the different layers is perturbed, which leads to abnormal contractions, the redistribution of the embryo incompressible content and the bursting of the yolk. The embryo to compensate ectopic stresses elongates or burst taking into account volume conservation.

Membrane removal, specifically macropinocytosis, has been found to associate to the folding or corrugation of cell surfaces. On a theoretical basis, high curvature, for a given transmembrane pressure, would reduce tension at the membrane and facilitate endocytosis to restore cell turgidity. Conversely, high tension would stall the endocytic machinery ^24^ Thus, cell membrane folds associate to surface contractile potential and membrane topology could be consider a good readout of surface stress and actomyosin activity. While membrane remodeling has been shown to be important in different morphogenetic events ^25–28^, its relation to actomyosin contraction has not been investigated in depth. We have found that during epiboly, surface stress and myosin accumulation at the yolk cell cortex display opposite profiles; stress is low in regions with high myosin (folded areas of the E-YSL) and high in regions with low myosin (smooth domains of the YCL) ^2^ Smooth membranes define regions under high tension while folded membranes associate to areas of active contraction. This consolidates the likelihood that the actomyosin contraction will generate membrane folds promoting endocytosis by lowering membrane tension. Myosin contraction as discussed above could eventually be prompted by mechanical cues or simply downstream of signaling events.

During epiboly, laser microsurgery in the yolk cell cortex leads to myosin recruitment and the formation of supernumerary membrane folds, which emphasizes the active character of membrane folding ^2^ Further, the observed spatial and temporal coordination of the stretching of the yolk cell membrane and the relaxation of the cortex underscores their intimate coupling. Accordingly, we propose that the natural membrane folds of the E-YSL would reflect the contraction of the underlying actomyosin cortex. Indeed, myosin distribution in the yolk surface ^29^ and membrane kinetics ^2^ at different epiboly stages follow the same regime.

### The DCs contribution

At early epiboly stages, DCs are positioned ahead of the EVL-E-YSL contact site, supporting the possibility that their radial intercalation could have an influence in the EVL vegetalward movement by shear contact. Indeed, the irregular distribution of the EVL cells at the margin and their rounded shapes in these early stages suggest that they are not the source of any pulling force. Further, the kinematics of the yolk granule’s flows, which parallels that of the blastoderm, E-YSL and EVL hint at a direct implication of DCs in this first epiboly stage. As described above, the DCs in direct contact with the yolk content would participate into the positive mechanical loop triggered at the E-YSL. After DC’s invagination, however, these cells lose contact with the E-YSL and a gap develops in between the two tissues, making their contribution to late epiboly progression unlikely. At this time, the DCs in contact with the I-YSL initiate retrograde movements (animalward), opposing those of the yolk granules.

### A diversity of mechanical models for Epiboly

Numerical simulations to mechanically model tissue expansion during epiboly (both cell shape changes and cell rearrangement) have been attempted from the early nineties ^30^. For *Fundulus* epiboly, the EVL was represented as a 2D sheet taking into consideration the balance between elastic forces and internal pressure (hydrostatic/osmotic pressure) within each cell. Modulating this balance would prompt cells to contract (higher elastic force) or to develop protrusive behavior (lower elastic tension than internal pressure). With these premises, cell behaviors were modeled such they would fit the experimental cell shape and rearrangements observed (Figure 5A and 5B). In particular, marginal EVL cells mechanical properties were tailored to display a net protrusive force at the margin with the yolk. In other words, local differences in tension, smaller at the boundary of the tissue than in between cells were described. In real terms, the tension topography was explained by either the active expansion of the EVL leading edge cells or by the pulling exerted by the E-YSL. From the margin, if the rate at which the mechanical stress dissipated through the epithelium was smaller than the strength of the applied external force, a gradient of stress would be predicted. This, indeed, seems to be the case in the EVL in both *Fundulus* and zebrafish (Figure 5C) ^7, 10^

**Figure 5.**
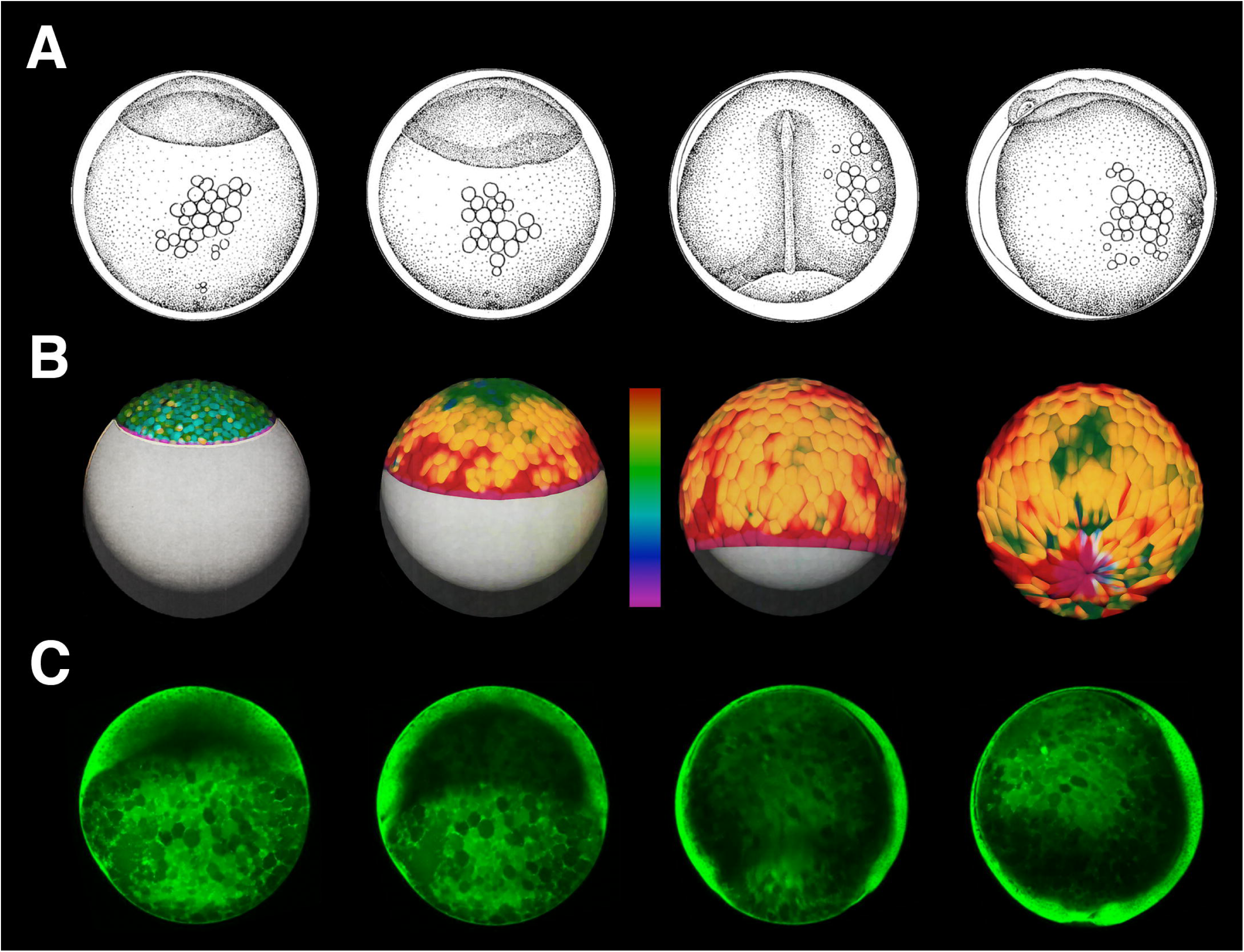
Numerical modeling of EVL elastic forces. **A)** Lateral view drawings of Fundulus embryos at: stage 14, epiboly has initiated and the blastoderm elevates at the animal pole; stage 15, the germ ring and the embryonic shield are visible; stage 19, dorsal view of an embryo at 70% epiboly presenting the convergent extension of the DCs; and stage 20, closure of the blastoderm around the yolk. Adapted from. **B)** Model simulation of EVL epiboly in Fundulus embryos. Different colors represent the balance of pressure (osmotic/hydrostatic) and circumferential elastic tension forces within each cell: green/blue when forces are equilibrated; red when cells are under elastic tension; and purple when nodes have protrusive behavior (higher pressure than elastic tension). Forces are calculated by the sum of forces at edge nodes. At epiboly onset most cells show low tension but gradually a gradient of stress develops from the EVL boundary towards the A pole. Cells closer to the yolk experience higher mechanical stress (red color) than cells near the A pole (green/blue color). Adapted from ^30^. **C)** Snapshots of a two-photon microscopy time-lapse movie at the medial plane of a membrane labeled transgenic zebrafish embryo [Tg (β-actin:mGFP)] at equivalent stages as Fundulus embryos in **A**.

The numerical simulations also predict that individual cells will acquire different shapes depending on their positions and the location of the applied pulling forces (originating at the EVL margin cells, at the E-YSL or from any other origin). Cells at different positions on the surface will experience different longitudinal (AV) and circumferential (CC) stress. At the A pole will be stretched equally in orthogonal directions, while near the margin they will suffer strong anisotropic stresses. Crossing the equator would also influence CC stresses, which will progressively relax with the reduction of the edge perimeter (the AV stress would not be affected). In summary, stress dissipation at each point would define the local cell shape anisotropies. Remarkably, as epiboly progresses, the contact area with the yolk margin of EVL cells is not uniform. While some of them keep wide contacts for a while, others exhibit elongated shapes and tend to be displaced out of the margin. These differences (EVL cells length/width ratio) correlate with distinct local levels of actin in the yolk ^11^. As a corollary, disparities in cells shapes appear to be a consequence of variations in mechanical stress. It is interesting to note that in Rab5ab MOs, where the E-YSL pulling force is reduced and epiboly is slowed down, elongation of cells in the AV axis at the margin was clearly reduced while they flatten along the circumferential axis ^20^. In this scenario as the balance of stresses is readjusted the stress dissipation in relation to the force applied would be higher.

These early numerical simulations quite accurately describe the behavior of cells at the surface within a tensional landscape. However they did not provide information about the sources or the molecular character of power and stress and did not take in consideration all the participating elements or the 3D distribution of these elements. Recently, a mechanical model considering the actomyosin belt in the E-YSL as the main source of power for epiboly progression has been proposed ^29^ This contractile belt would just work after the EVL surpasses the equator bringing together all tissues at the V pole. CC contraction would be coupled to the embryo geometry and AV contractility would be negligible. Yet, CC contractility would not suffice to justify AV progression before the EVL reaches the equator, neither will explain the AV tensional patterns detected by laser cuts. Thus, a complementary element, a flow-friction motor, was claimed. This would generate the additional AV tension necessary for progression. The flow-friction force would originate in the E-YSL ring from actin and myosin retrograde cortical flows (Figure 6A). These flows would have different speed than the adjacent layers displacements (the yolk cell membrane, the cytoplasm or the microtubules network) and friction will be generated. Indeed, it was determined that microtubules move at a different speed than the actomyosin microfilaments and the cytoplasm and yolk cell membrane do not move at all ^29^, although no formal prove of friction was presented.

**Figure 6.**
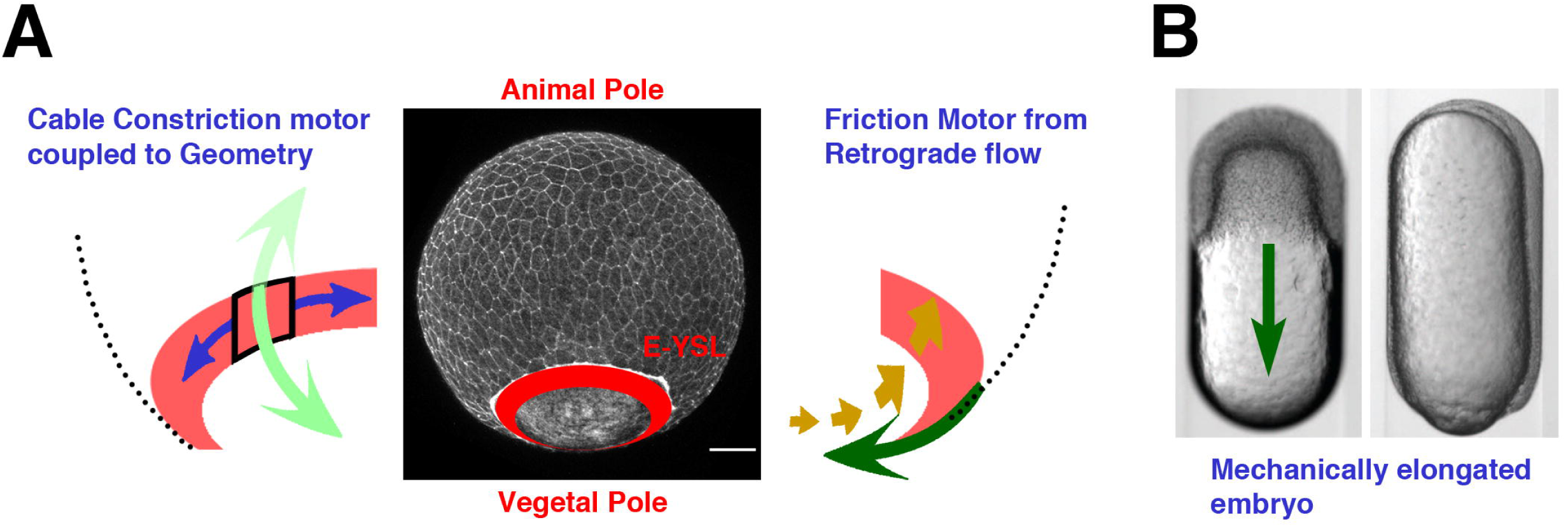
The flow-friction model. **A)** Schematic representation of the theoretical model. The EVL and the actomyosin ring were modeled as two thin compressible viscous fluids mechanically connected generating internal active tension. The circumferential tension at the contractile E-YSL (left) will move the ring to the nearest pole, while actomyosin retrograde cortical flows coupled to friction against the underlying substrate (right) will generate a pulling force in the EVL directed to V. **B)** Bright field images of an embryo inside a cylindrical agarose tube at 30% epiboly (top) and at closure (bottom). A green arrow indicates epiboly progression. Adapted from ^29^ Both, the flow-friction and the differential tension models are fully compatible with epiboly in elongated embryos.

The actomyosin ring and the EVL were modeled as compressible isotropic thin-film fluids, the shear and bulk viscosities were accounted and an additional active tension generated by the actomyosin network (contraction) was considered. As a boundary condition it was assumed that the ring had a mechanically free interface as its V side. This assumption was supported by the absence of cortex recoil at the V pole after laser microsurgery and by the observed exponential decrease of myosin intensity at the V side of the contractile E-YSL ring. This was understood to correlate with an exponential decrease of active tension within the ring in the V direction (not proved either). The active tension profile was approximated to the myosin intensity distribution. Finally, the ring width was estimated from experimental measurements of actin and myosin. This theoretical model fits the experimental cortical flow profiles, the ring velocity during epiboly and the relative tensions (CC and AV) in the ring extracted by laser cuts.

Experimentally distorted AV-cylindrically shaped embryos that manage to undergo epiboly have been claimed to support the flow friction model ^29^ In these embryos, the directional effect of the CC tension generated at the actomyosin E-YSL ring is cancelled by the geometry. Still, epiboly progresses vegetalwards, which makes the CC contraction of the ring dispensable (Figure 6B). If a flow-friction force is discarded, a pure CC contractile ring could not explain the experimental tension patterns, neither could make epiboly progress in a cylindrical conformation.

Yet, the flow-friction model does not succeed in explaining in global terms how epiboly proceeds. The E-YSL contractile ring is in fact not mechanically isolated from the rest of the yolk cell as it was assumed. The EVL, the E-YSL and the YCL are all in a *continuum.* AFM analyses demonstrate that the yolk cell surface behaves, at the time scale of interest, as elastic, but more importantly show that there is tension at the V pole of the yolk cell ^2^. Although this tension may be passive (tensile), it should still constitute a mechanical link. Further, the friction mechanism fails to clarify the disappearance of the yolk cell membrane, which is essential for epiboly progression ^20^, or the progressive recruitment of the YCL in the E-YSL. A second argument against the model comes from the detailed observation of the cortical flows.

Actin and myosin retrograde flows take place at all times outside the ring or folded area (E-YSL), while in the theoretical model the resultant friction generates and propagates from inside the E-YSL ^29^

Indeed, the differential tension / membrane removal model emanating from the HR analyses ^2^ is also fully compatible with epiboly in an elongated embryo. As the orthogonal actomyosin contraction (AV and CC) at the E-YSL acts in the 2D surface plane, the embryo geometry, spherical or elliptical, will make no much difference. The differences in the mechanical properties of the adjacent domains (EVL and YCL) and yolk cell membrane endocytosis will still perform in cylindrical embryos and likewise epiboly will progress.

We consider that the differential tension model manages to explain all data inferred and experimental, the dynamic mechanical equilibrium of the whole embryo, the contribution of its elastic surface, the yolk cell membrane removal and the yolk internal flows dynamics. It globally elucidates the full process of epiboly (Figure 3B).

## EXPERIMENTAL PROCEDURES

### Zebrafish lines maintenance

AB and TL wild type strains were used throughout this study. Membrane-GFP transgenic (Tg (β*-actin:m-GFP*)) fish ^31^ were provided by Lilianna Solnica-Krezel. Adult fish were cultured under standard conditions and staged embryos were maintained at 28.5 °C in embryo medium ^32^

### Morpholino injections

To knockdown Rab5ab, morpholino yolk injections (4 ng and 8 ng) were performed at the 512-1024-cell stage. Morpholinos (MOs) were purchased from Gene Tools and designed against selected regions (ATG or UTR) of the *rab5ab* gene (Accession Number ENSDARG00000007257) (Gene Tools):

MO1-ATG (5-TCGTTGCTCCACCT-CTTCCTGCCAT-3)

MO2-UTR (5-GACCCAAAACCCCAATCTCCTGTAC-3)

MO3-UTR-ATG (5-ACCTCTTCCTGCCATGACCCAAAAC-3)

Mismatch MO (5-TCcTTcCTCgACCTCTTCgTcCCAT-3) (mispaired nucleotides in lower case) ^20^.

For all experiments, a group of embryos was injected with a Standard Control MO (5-CCTCTTACCTCAGTTACAATTTATA-3).

### Live Imaging and Analysis

Whole embryo images were collected from non-dechorionated animals aligned in a 1.2 % agarose mold and covered by E3 medium. Images were acquired (4X magnification) every 5 minutes with an Olympus MVX10 Macroscope.

For confocal microscopy, embryos were mounted in 0.5 % low melting agarose (A9045 Sigma) in E3 embryo medium.

Sagittal sections (350 μm depth from the yolk cell membrane surface) were collected from (Tg (*β-actin: m-GFP*)) embryos using a Leica SP5 two-photon microscope equipped with a mode-locked near-infrared MAITAI Laser (Spectra-Physics) tuned at 900 nm, with non-descanned detectors and with a 25 × / 0.95 water-dipping objective. Images were scanned at 200 Hz and frames were averaged three times. Stacks of 30 μm, 10 μm step-size, were acquired every 2 minutes.

Most image analyses and processing were performed using Fiji (http://pacific.mpi-cbg.de) and Matlab (Mathworks).

## ACKNOWLEDGEMENTS

We thank the Confocal Microscopy Unit from IBMB-PCB, the Advanced Digital Microscopy Core Facility from IRB Barcelona, Xavier Esteban and members of the laboratory for continuous support. We are grateful to Amayra Herna ández-Vega for sharing data and thoughts and Carolina Minguilloón for reading earlier versions of this manuscript. The Consolidated Groups Program of the Generalitat de Catalunya and DGI and Consolider Grants from the Ministry of Economy and Competitivity of Spain to EMB supported this work.

## AUTHOR CONTRIBUTION

MM performed all biological tests; EMB designed the study, analyzed the data and wrote the paper. All authors discussed the results and commented on the manuscript.

